# Tracing the genetics of neurological disease to the mutation-directed addition of single hydrogen bonds

**DOI:** 10.1101/2021.12.05.471334

**Authors:** Xiaoming Zhou, Kyuto Tashiro, Lily Sumrow, Lillian Sutherland, Glen Liszczak, Steven L. McKnight

**Affiliations:** Department of Biochemistry, UT southwestern Medical Center, Dallas, TX 75390, USA

## Abstract

Mutations causative of neurological and neurodegenerative disease can occur in coding regions that specify protein domains of low sequence complexity. These autosomal dominant mutations can be idiosyncratic in their recurrent appearance at the same amino acid. Here we report studies of recurrent mutations in proline residues located within low complexity (LC) domains associated with the neurofilament light chain protein, the microtubule-associated tau protein, and the heterogeneous nuclear RNPA2 protein. All such mutations manifest their effects by directing formation of variant proteins endowed with the addition of a single, main chain hydrogen bond specified by the variant amino acid replacing proline. Here we show that methylation of the peptide backbone nitrogen atom associated with these variant amino acids eliminates the aberrant hydrogen bond and restores normal protein function.

## Introduction

Human genetic studies represent a powerful approach to the understanding of disease. Upon completion of the human genome, human genetics began to enter its golden age. Over the past several decades the field of human genetics has opened the door to an enhanced molecular understanding of innumerable diseases. In an ever-growing number of cases, this newfound knowledge has created the opportunity for rational therapeutic intervention.

Maturation of the knowledge created by genetic studies is invariably assisted by biochemical and biological research. An understanding of the functional role of proteins altered by disease-causing mutations creates both satisfaction associated with the logic of new scientific truth, as well as the opportunity to consider rational avenues for therapeutic intervention.

The translational pathway evolving from genetic studies of neurological and neurodegenerative disease has been impeded at the bridge connecting genetics to biochemistry. Human mutations causative of neurological or neurodegenerative disease are often represented by missense mutations that fall in protein domains of low sequence complexity. These low complexity (LC) domains have long been believed to function in the absence of any form of structural order (*1*, *2*). As such, it has been difficult to understand the functional consequences of subtle, missense mutations. How might such variants have any functional impact upon a protein domain if it is devoid of structure?

Here we report studies on LC domains associated with the neurofilament (NFL) light chain protein, the microtubule-associated protein tau, and the heterogeneous nuclear RNPA2 (hnRNPA2) protein. Each of these proteins is known to be subject to autosomal dominant mutations in proline residues causative of disease. Our observations provide a biochemical description of how these recurrent, idiosyncratic mutations affect protein function.

## Results

Human Charcot-Marie-Tooth (CMT) disease is caused by mutation of either of two proline residues located within the head domain of the neurofilament (NFL) light chain protein (*3*, *4*).These autosomal dominant mutations change proline residue 8 to leucine, arginine or glutamine, and proline residue 22 to serine, threonine or arginine. Various assays have been used to confirm that these mutations impede the proper assembly of neurofilaments in living cells (*3*, *5*, *6*).

Intermediate filament (IF) proteins can be expressed in bacterial cells, purified and incubated under conditions permissive of formation of mature filaments. The first two steps in assembly - dimerization and tetramerization - are mediated solely via centrally located, α helical rod domains and can proceed in the absence of the head domain. By contrast, the head domains of IF proteins are essential for the subsequent assembly of eight tetramers into cylindrical filaments (*7*–*9*).

The head domains of intermediate filaments are of low sequence complexity and have long been thought to function in the absence of molecular structure (*10*, *11*). An isolated report published five years ago offered the contradictory claim that IF head domains adopt *bona fide* molecular structure so as to facilitate self-association in a manner resulting in phase separation (*12*). In this regard, IF head domains share functional relatedness with the LC domains of certain RNA binding proteins that self-associate in the form of liquid-like droplets and hydrogels (*13*–*15*).With the help of valuable collaborators, we published a follow-up report focused on the head domains of the desmin and NFL intermediate filament proteins earlier this year (*16*). Both head domains, in isolation, were shown to form labile cross-β structures as characterized by solid state NMR spectroscopy. By use of intein ligation enabling segmental isotopic labeling, it was possible to show that the NMR spectra of the isolated desmin and NFL head domains match the spectra observed in fully assembled intermediate filaments.

Such studies rejected the notion that IF head domains function in the absence of molecular structure, and instead concluded that they form labile and reversible cross-β structures. This concept of LC domain self-association conforms with similar studies on LC domains associated with RNA binding proteins, including hnRNPA2, yeast ataxin-2, fused-in-sarcoma, and TDP-43 (*13*–*15*, *17*–*20*).

The recent Zhou et al. (2021) study of IF head domains also evaluated human mutations in desmin causative of cardiac deficits, and NFL causative of CMT disease (*16*). All desmin and NFL head domain mutations were found to enhance the stability of otherwise labile cross-β interactions. Such observations comported with the autosomal dominant nature of the human mutations in desmin and NFL, and offered the idea that aberrantly enhanced strength of head domain self-association is incompatible with proper intermediate filament assembly.

The proximity of the P8 and P22 CMT mutations to the observed position of labile cross-β structure within the NFL head domain is reminiscent of findings on the TDP-43 RNA binding protein described in the accompanying paper (Zhou et al., manuscript submitted). Since proline is the only amino acid incapable of participating in Pauling hydrogen bonding, we reasoned that proline residues 8 and 22 within the NFL head domain might locally insulate against the aberrant spread of cross-β structure.

When either proline residue 8 or 22 of the NFL head domain is changed to any of the other nineteen amino acids, the capacity for Pauling hydrogen bonding is universally restored. This restoration might allow for the spread of cross-β structure, resulting in pathologic strengthening of head domain self-association. This idea was tested via methyl capping of the peptide backbone nitrogen atoms associated with the leucine, arginine and glutamine variants of P8, as well as the serine, threonine and arginine variants of P22.

A native chemical ligation reaction was used to link synthetic peptides containing single, N^α^-methyl amino acids to the remaining 517 residues of the NFL polypeptide (Fig. 1A). These semi-synthetic NFL proteins were purified and incubated under conditions receptive to assembly of mature intermediate filaments. As shown in Fig. 1B, NFL proteins bearing any of the six CMT mutations formed amorphous tangles. By contrast, all six of the methyl cap-repaired variants formed morphologically normal intermediate filaments indistinguishable from those assembled from the native NFL protein.

**Fig. 1.**
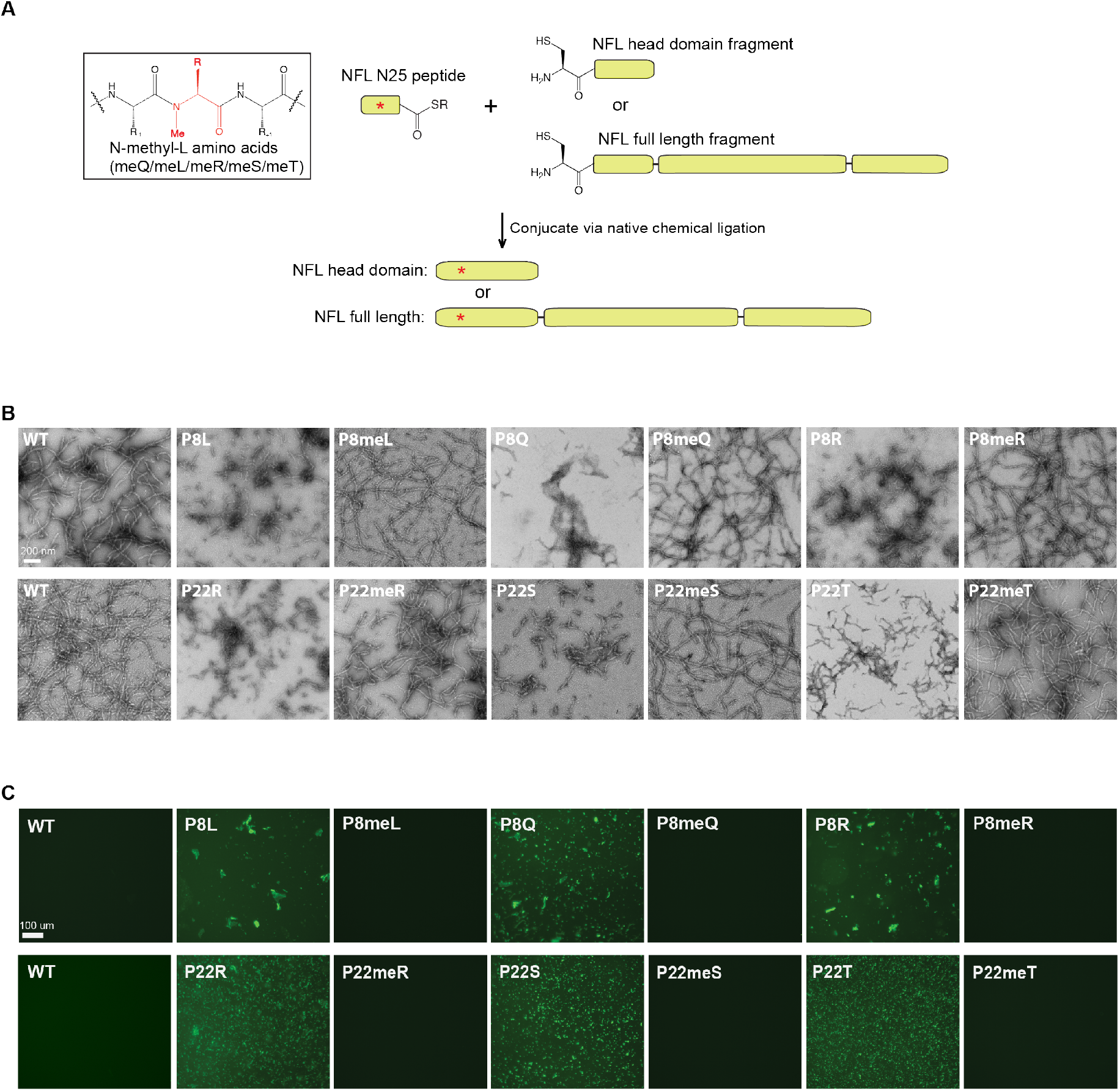
Effects of methyl-capping peptide backbone nitrogen atoms associated with disease-causing residues within the neurofilament light (NFL) chain head domain. Synthetic peptide thioesters corresponding to the amino-terminal twenty five residues of the NFL protein were prepared to contain: (i) the native sequence of NFL; (ii) Charcot-Tooth-Marie (CMT) mutational variants at position 8 or 22; or (iii)CMT variants methyl-capped at the peptide backbone nitrogen in the mutated amino acid. Native chemical ligation was used to produce full length NFL protein or the isolated head domain bearing different synthetic peptides at the aminoterminus **(Panel A)**. Detailed procedures employed to construct and purify semi-synthetic NFL derivatives are described in Materials and Methods. Full length proteins carrying P8L, P8Q and P8R variations, P22R, P22S or P22T variations, or N^α^-methyl variants of these six residues, were incubated under conditions receptive to the formation of intermediate filaments (Materials and Methods). Filament assembly was monitored by transmission electron microscopy **(Panel B)**. Assembly assays for all six CMT variants yielded tangled, amorphous precipitates. Assembly assays for semi-synthetic proteins containing a methyl-capped nitrogen of the individual CMT variant amino acid residues (the L, Q and R variants of position 8, and the R, S and T variants of position 22) yielded homogenous intermediate filaments indistinguishable from those made from the native NFL protein. Thioflavin-T staining of synthetic peptides used to generate semisynthetic, full-length NFL proteins revealed dye-stained precipitates for each peptide bearing a CMT-causing lesion (P8L, P8Q, P8R, P22R, P22S and P22T). No dye-stained precipitates were observed for the parental peptide (WT), and no dye-stained precipitates were observed for the CMT-causing variants that were modified to methyl-cap the peptide backbone nitrogen atom of the variant amino acid residue **(Panel C)**. Scale bar in Panel B = 200 nm, Panel C = 100 μm.

In the process of building and purifying the peptides employed for production of the semi-synthetic NFL variants used in this study, it was noticed that the P8Q, P8L, P8R, P22R, P22S and P22T peptides were more prone to precipitation than the peptide bearing the native sequence. More surprisingly, the methyl-capped versions of these six CMT-mutated variants were uniformly more soluble than the parental substrates bearing normal peptide nitrogen atoms at the P8Q, P8L, P8R, P22R, P22S and P22T positions.

Purified peptide samples containing the N-terminal twenty five residues of NFL, and mutant variants thereof, were mixed with thioflavin-T, spotted onto microscope slides and visualized by fluorescence microscopy (Fig. 1C). The peptide containing the native NFL sequence showed no evidence of precipitation and thioflavin-T staining. The peptides containing each of the six CMT-causing variations in the NFL head domain all showed prominent, thioflavin-T stained precipitates. In all six cases, methyl-capping of the peptide backbone nitrogen atom associated with the variant amino acids fully solubilized the head domain peptides.

Enhanced peptide precipitation (Fig. 1C) causative of aberrant IF assembly (Fig. 1B) was assumed to result from mutation-directed strengthening of NFL head domain self-association. To test this prediction, synthetic peptides carrying the P8Q and P22S mutations, or N^α^-methyl derivatives thereof, were ligated to the remainder of the NFL head domain (Fig. 1A). Ligation products were purified and incubated under conditions known to distinguish between the solubility of the native head domain of NFL relative to CMT mutants (*16*). As shown in Supplemental Data Fig. S2, the native NFL head domain was soluble under such conditions, variants carrying either the P8Q or P22S mutations were not, and methyl-capping of the peptide backbone nitrogen atoms associated with P8Q and P22S fully restored solubility.

We reason that each of the three variants at the P8 position, as well as the three variants at the P22 position, cause the untoward strengthening of an otherwise labile and transient cross-β structure useful for NFL head domain self-assembly. Enhanced strength of self-association is interpreted to result from an extended hydrogen bond network, and this extension is prevented by methyl capping of the peptide backbone nitrogen atoms associated with the variant amino acid residues. To further probe this concept we employed a different chemical approach to deny hydrogen bonding by one of the variant amino acid residues. Instead of methyl capping the peptide backbone nitrogen associated with the leucine residue of the P8L CMT mutant, we replaced the peptide backbone nitrogen with oxygen (Fig. 2A).

**Fig. 2.**
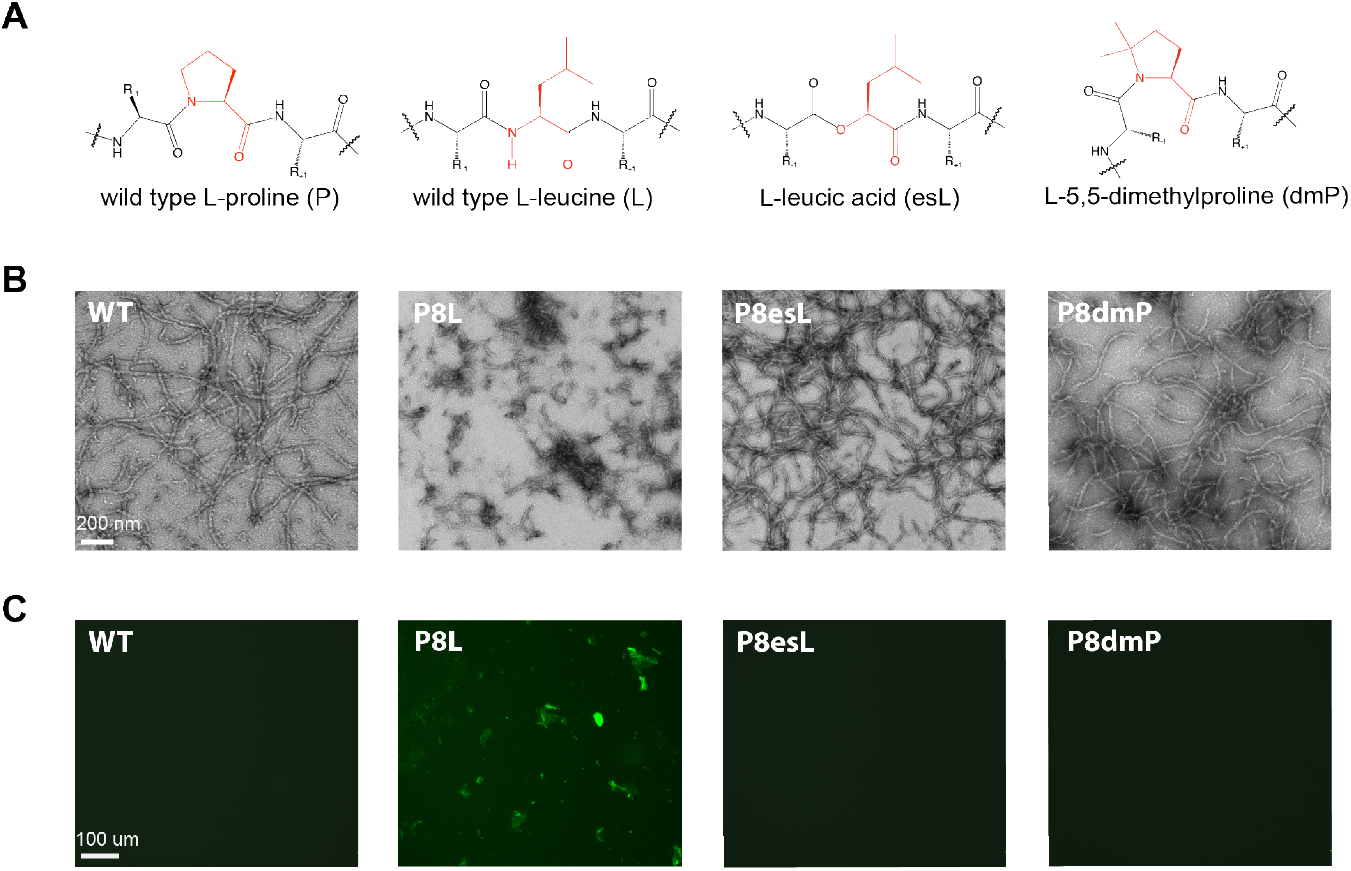
Effects of eliminating Pauling hydrogen bonding via amide-to-ester substitutions in the peptide backbone, and effects of replacing proline 8 of the NFL head domain with dmP. Synthetic peptides were prepared replacing proline residue 8 of the NFL head domain with leucine, L-leucic acid or dmP. **Panel A** shows the structures of synthetic peptides with chemical variation highlighted in red. Semi-synthetic, full length NFL proteins bearing the structures shown in Panel A were assembled, purified and assayed under conditions receptive to the assembly of mature intermediate filaments **(Panel B)**. Reconstruction of the P8L CMT variant yielded tangled, amorphous precipitates. Amide-to-ester backbone substitution at leucine residue 8 (P8esL), the eliminating a hydrogen bond donor from this site, facilitated assembly of intermediate filaments indistinguishable from those assembled from the native NFL protein. Replacement of proline residue 8 with dmP (P8dmP) allowed assembly of homogeneous intermediate filaments morphologically identical to those produced by the native protein (WT). Thioflavin-T staining of synthetic peptides used to generate semi-synthetic, full length NFL proteins revealed dye-stained precipitates in the case of the peptide bearing the P8L variant **(Panel C)**. Esterification of the peptide backbone at leucine residue 8 fully solubilized this peptide. Replacement of proline residue 8 with dmP yielded a peptide having solubility properties indistinguishable from the peptide bearing the native sequence of NFL (WT). Scale bar in Panel B = 200 nm, Panel C = 100 μm.

A synthetic peptide bearing an amide-to-ester backbone substitution at the P8L site was ligated to the remainder of the NFL protein. The semi-synthetic protein was purified and incubated under conditions suitable for assembly of intermediate filaments. As shown in Fig. 2B, this amide-to-ester variant formed intermediate filaments indistinguishable from those assembled from the native NFL protein. In other words, esterification at the appropriate position of the polypeptide chain repaired the P8L deficit to an extent indistinguishable from methyl-capping of the P8L peptide backbone nitrogen. We further observed that the amide-to-ester substitution fully solubilized the P8L synthetic peptide as determined by thioflavin-T staining and fluorescence microscopy (Fig. 2C).

Other than being the sole amino acid incapable of forming Pauling hydrogen bonds, proline is also unique in the flexibility it allows for rotation of the peptide backbone relative to the other nineteen amino acids. As a means of probing the potential importance of this chemical feature of proline, we replaced proline residue 8 of the NFL head domain with L-5,5-dimethylproline (dmP). Relative to the normal proline side chain, which strongly prefers the *trans* conformation, dmP is strongly disposed towards the *cis* isomeric conformation (*21*). A synthetic peptide carrying dmP at residue eight was ligated to the remainder of NFL. The purified ligation product was tested for capacity of the semi-synthetic protein to assemble into intermediate filaments. As shown in Fig. 2B, the dmP derivative assembled into intermediate filaments indistinguishable from those formed from the native NFL protein. It was likewise observed that the dmP-containing peptide was equally soluble to the native peptide corresponding to the first twenty five residues of the NFL head domain as assayed by thioflavin-T staining and fluorescence microscopy (Fig. 2C). These experiments provide no evidence that phenotypic deficits exacted by NFL proline mutations result from impediments to rotation of the peptide bond.

Having begun to understand why the mutation of NFL to any of a number of other amino acids in place of proline 8 or proline 22 manifests in a biochemical sense, we looked for similarly idiosyncratic proline mutations causative of neurodegenerative disease. Perhaps the most penetrant mutation of the cytoplasmic tau protein causative of neurodegenerative disease maps to proline residue 301. Disease causing changes of this P301 residue include substitution by leucine, serine or threonine(*22*–*28*). Knowing that tau mutations prompt formation of amyloid-like aggregates (*29*, *30*), we reasoned that P301 might serve the normal protein in a manner insulating against the spread of localized cross-β structure.

Previous studies have already provided compelling evidence that synthetic peptides corresponding to residues 295–311 of tau suffer profound propensity to aggregate if proline residue 301 is changed to either leucine or serine (*30*). The importance of peptide backbone rotation was favored according to the differential effects of substituting proline with either the *cis* (2S,4S) or *trans* (2S,4R) conformer of 4-fluoroproline (Fig. 3A) (*31*). The latter conformer was reported to favor peptide aggregation in a manner similar to the P301L and P301S mutations.

**Fig. 3.**
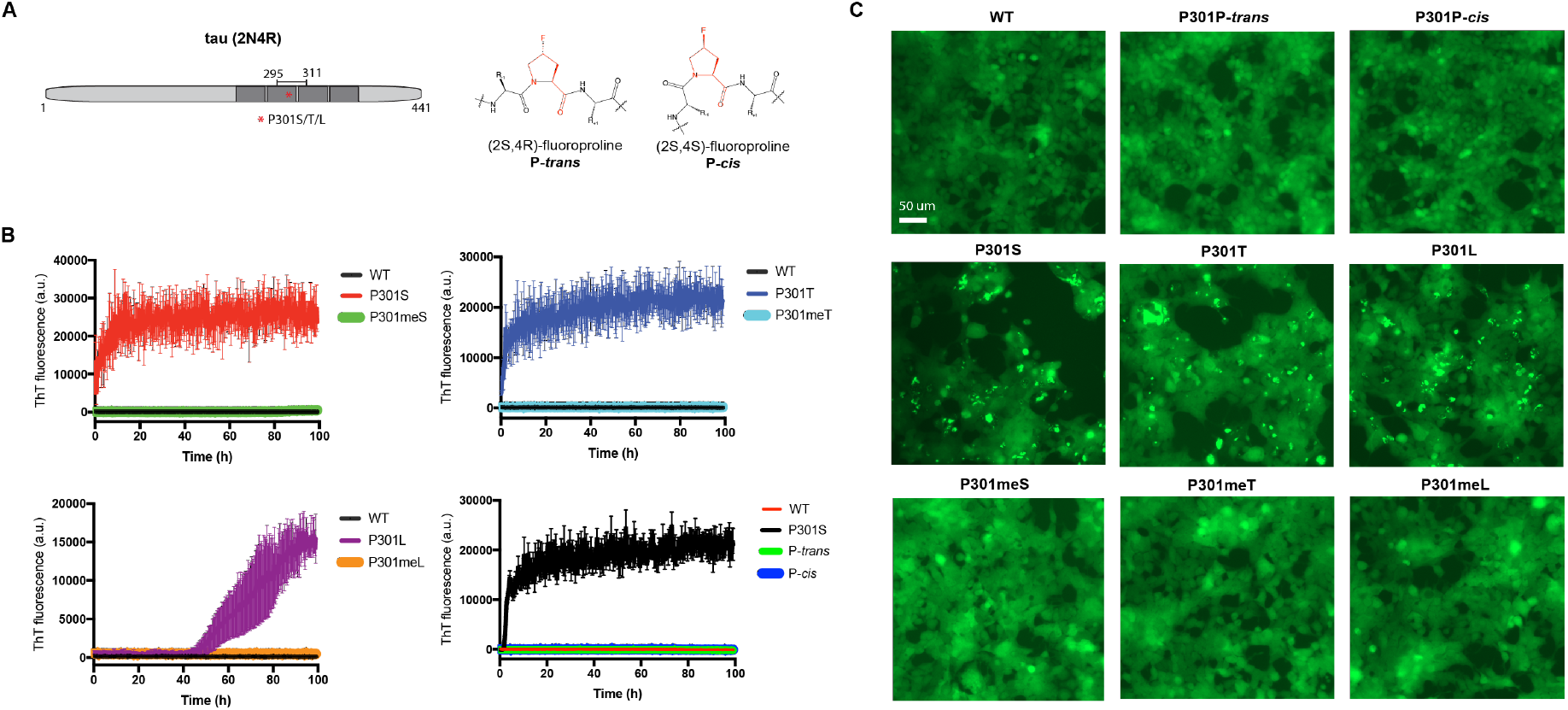
Restorative effects of eliminating Pauling hydrogen bonding by methyl-capping of peptide backbone nitrogen atoms of the P301S, P301T and P301L mutational variants of tau. Peptides corresponding to residues 295–311 of the cytoplasmic tau protein were synthesized in the native form of the protein and in the forms of three disease causing variants (P301S, P301T and P301L). Additional derivatives included methyl-capped forms of the three disease variants, as well as variants of the native peptide wherein proline residue 301 was replaced by either the *trans* (2S,4R; *P-trans*) or *cis* (2S,4S; P-*cis*) conformer of 4-fluoroproline. The locations of the variants within the intact tau polypeptide, and chemical structures of the *P-trans* and P-*cis* variants, are shown in **Panel A**. Each sample was mixed with thioflavin-T, incubated and monitored for the appearance of enhanced fluorescence **(Panel B)**. Each of the disease-causing variants led to strong increases in thioflavin-T fluorescence relative to the peptide bearing the native sequence of tau. No such increase was observed if the peptide backbone nitrogen atoms associated with the disease variant residues were methyl-capped. No enhancement of thioflavin-T fluorescence was observed for peptides wherein proline residue 301 was replaced by either the *P-trans* or P-*cis* conformer of 4-fluoroproline. Each of the eight synthetic peptides were subjected to lipofection-mediated transfer into a tau biosensor cell line expressing a fluorescent tau fusion protein. No change in the homogenous, wide-spread distribution of endogenous fluorescent tau was observed in cells transfected with either the native tau peptide or peptides wherein proline residue 301 was replaced by either conformer of 4-fluoroproline **(Panel C)**. Each of the peptides bearing a disease-causing substitution led to the formation of distinct aggregates upon assay in the tau biosensor cell line. When the peptide nitrogen atoms associated with the disease variant residues (serine, threonine or leucine) were methyl-capped, the resulting distribution of endogenous tau in the biosensor cells was equivalently homogenous and wides-pread as compared with the synthetic peptide specifying the native tau sequence. Scale bar = 50 μm.

We prepared these same peptides and confirmed that the P301L, P301S and P301T mutations profoundly affected solubility, relative to the peptide bearing the native tau sequence, as deduced by thioflavin-T fluorescence. We failed, however, to observe any difference in peptide solubility when either the *cis* or *trans* conformer of 4-fluoroproline was used in place of proline at residue 301 (Fig. 3B).

Extending from these observations we prepared peptide derivatives in which a methyl cap was placed on the peptide backbone nitrogen atom of the leucine, serine or threonine mutational variant residues. As shown in Fig. 3B, the methyl-capped variants of all three tau mutant peptides regained solubility indistinguishable from the native tau peptide. In other words, methyl capping fully repaired the deficits in peptide aggregation specified by the three tau mutations.

Moving from test tube reactions to living cells, we employed a tau biosensor cell line expressing tau-CFP and tau-YFP. The fluorescent tau protein in these cells is homogeneously distributed, yet can be converted to a punctate distribution when the cells are exposed to pathological tau aggregates (*32*, *33*). Each of the tau peptides studied in biochemical assays shown in Fig. 3 was internalized by lipofection into the tau biosensor cell line. The native tau peptide did not affect the intracellular distribution of fluorescent tau, nor did peptide variants carrying either conformer of 4-fluroproline. Peptides corresponding to the P301L, P301S and P301T mutations, upon introduction into the tau biosensor cells, led to obvious aggregation of the endogenous tau protein. Finally, synthetic peptide variants of the P301L, P301S and P301T mutants bearing a methyl cap on the leucine, serine or threonine residue caused no detectable aggregation of endogenous tau.

As a final example of human genetic studies unfulfilled because of shortcomings to our biochemical understanding of LC domains, we were attracted the familial, early-onset Paget’s disease mutation of proline residue 298 of the gene encoding hnRNPA2 (*34*). Our attention to this mutation, aside from its altering of a conserved proline residue, came from the fact that the mutation is located within an LC domain long understood to self-associate in a manner leading to phase separation. Indeed, the P298L Paget’s disease mutation is located in the middle of the labile cross-β core of the hnRNPA2 protein as characterized by biochemical footprinting, solid state NMR spectroscopy and cryo-electron microscopy (*14*, *19*, *35*).

A three-piece native chemical ligation strategy was implemented to incorporate a synthetic peptide encompassing the site of the P298L mutation into the remainder of the hnRNPA2 LC domain (Fig. 4A). Reconstruction of the native hnRNPA2 LC domain by these methods yielded a protein that phase separated into spherical, liquid-like droplets. The reconstructed variant carrying the P298L mutation formed distinctly misshapen droplets. Methyl-capping of the peptide backbone nitrogen atom associated with the variant leucine residue, followed by sequential ligations to reconstruct the intact LC domain, yielded a protein that phase separated into spherical, liquid-like droplets indistinguishable from those made by the native hnRNPA2 LC domain (Fig. 4B). As in the cases of the other nine mutations evaluated in this study, denial of the capacity for Pauling hydrogen bonding for the 298L residue fully rescued properly balanced self-associative capacity to the hnRNPA2 LC domain.

**Fig. 4.**
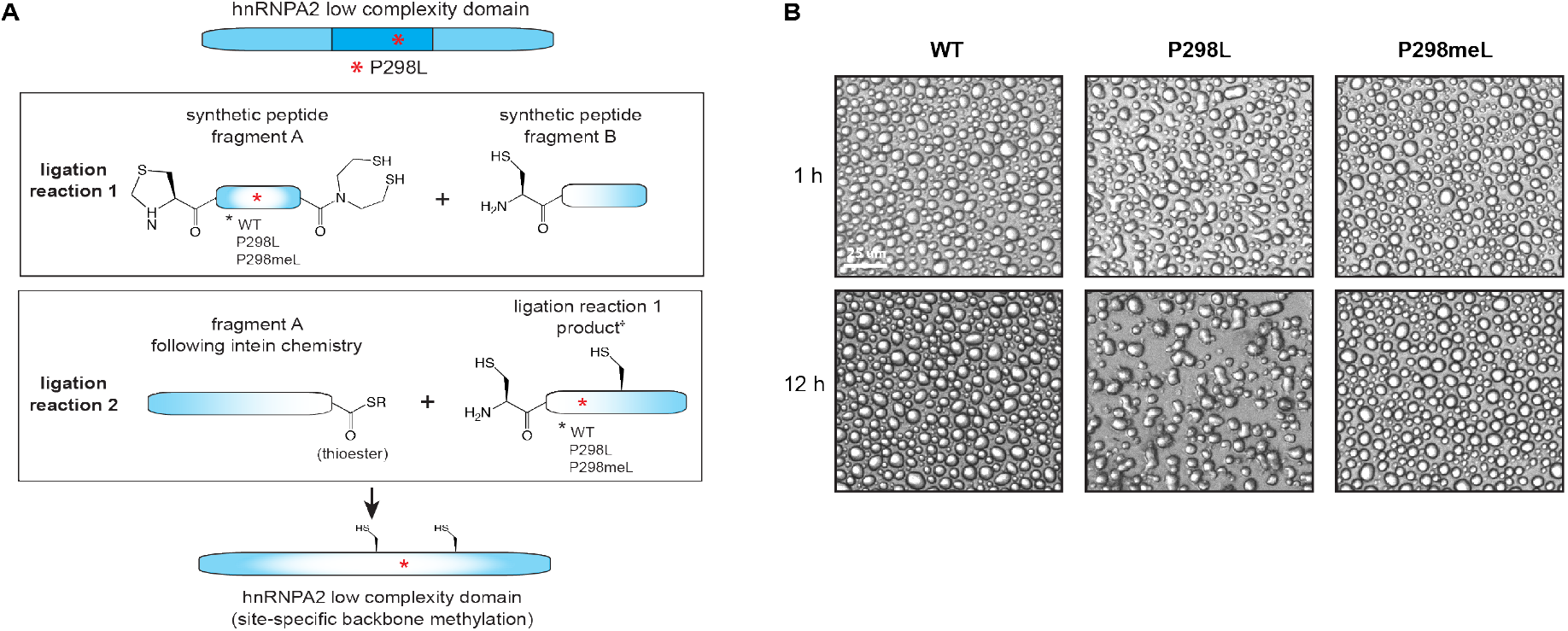
Restorative effect of eliminating the Pauling hydrogen bond by methyl-capping the peptide backbone nitrogen atom of the P298L mutational variant of hnRNPA2. A three-piece native chemical ligation strategy was used to prepare semi-synthetic variants of the fulllength low complexity (LC) domain of hnRNPA2 **(Panel A)**. Detailed methods for generation of the semi-synthetic hnRNPA2 LC domain are presented in Materials and Methods. The assembled hnRNPA2 LC domain contained either the native sequence, the disease-causing P298L variant, or a variant in which the peptide backbone nitrogen atom associated with the leucine 298 residue was methyl-capped. Following purification, the three samples were tested for phase separation in an aqueous buffer **(Panel B)**. The native LC domain of hnRNPA2 formed spherical, liquid-like droplets (left). The P298L variant formed distinctly misshapen droplets (middle). Methyl-capping of the peptide backbone nitrogen associated with leucine 298 residue yielded a semi-synthetic LC domain variant that formed spherical droplets indistinguishable from those formed by the native protein (right). Scale bar = 25 μm. ^**†**^ Ligation reaction 1 product shown after thiazoldine deprotection step.

## Discussion

In this study we report experiments focused on ten unusual mutations causative of human disease. All ten mutations localize to one of four proline residues, two in the head domain of NFL, one in tau, and one in hnRNPA2. Functional assays of the NFL, tau and hnRNPA2 proteins allowed us to probe how mutations of these four proline residues might manifest at a mechanistic level. In all of ten cases, we were able to functionally repair the mutations by eliminating the capacity of the variant amino acid residue to participate in Pauling hydrogen bonding.

We offer that this science is of value in two ways. First, it helps bridge human genetics and biochemistry. If our interpretations are correct, we are now able to understand how certain proline mutations function to change the properties of the LC domains in which they reside. The mechanistic understanding of how human mutations alter the properties of their encoded proteins should favor the opportunity for considering rational avenues for therapeutic intervention. Second, our experiments will facilitate bioinformatic searches for similarly idiosyncratic proline mutations described by geneticists over the past several decades. We predict that if a proline residue is the locus of recurrent mutations that change it to two or more different amino acid residues, and if the mutation lies within an LC domain, then the mechanistic basis for its deleterious effects might match what we describe for mutations in NFL, tau and hnRNPA2.

We close by warning readers that the scientific findings reported in this and the accompanying manuscript are inconsistent with the prevailing model for LC domain self-association currently accepted by the fast-paced “phase separation” field. For more than a decade we have reported experimental observations consistent with the interpretation that LC domains undergo self-association and consequential phase separation via the formation of labile cross-β interactions. These assemblies are heavily reliant on Pauling hydrogen bonding exacted by the main chain of polypeptides, and are reflective of structural interactions that are *bona fide* and specific. Almost all scientists working in the phase separation field have rejected our science and instead believe that LC domains function in the complete absence of protein structure. We hope that the data presented in this and the accompanying paper will help accelerate resolution between these conflicting perspectives.

## Acknowledgments

We thank Masato Kato, Deepak Nijhawan and Robert Thompson for technical advice and encouragement. This work was supported by an anonymous donor and National Institute of General Medical Science of NIH grant R35 GM130358 and National Cancer Institute of NIH grant U54 CA231649 to SLM. This work was also supported by Welch Foundation grant I-2039-20200401 and the Cancer Prevention Research Institute of Texas grant RR180051 to GL.

## Competing interests

Authors declare that they have no competing interests.

## Data and materials availability

All data are available in the main text or the supplementary materials.

---

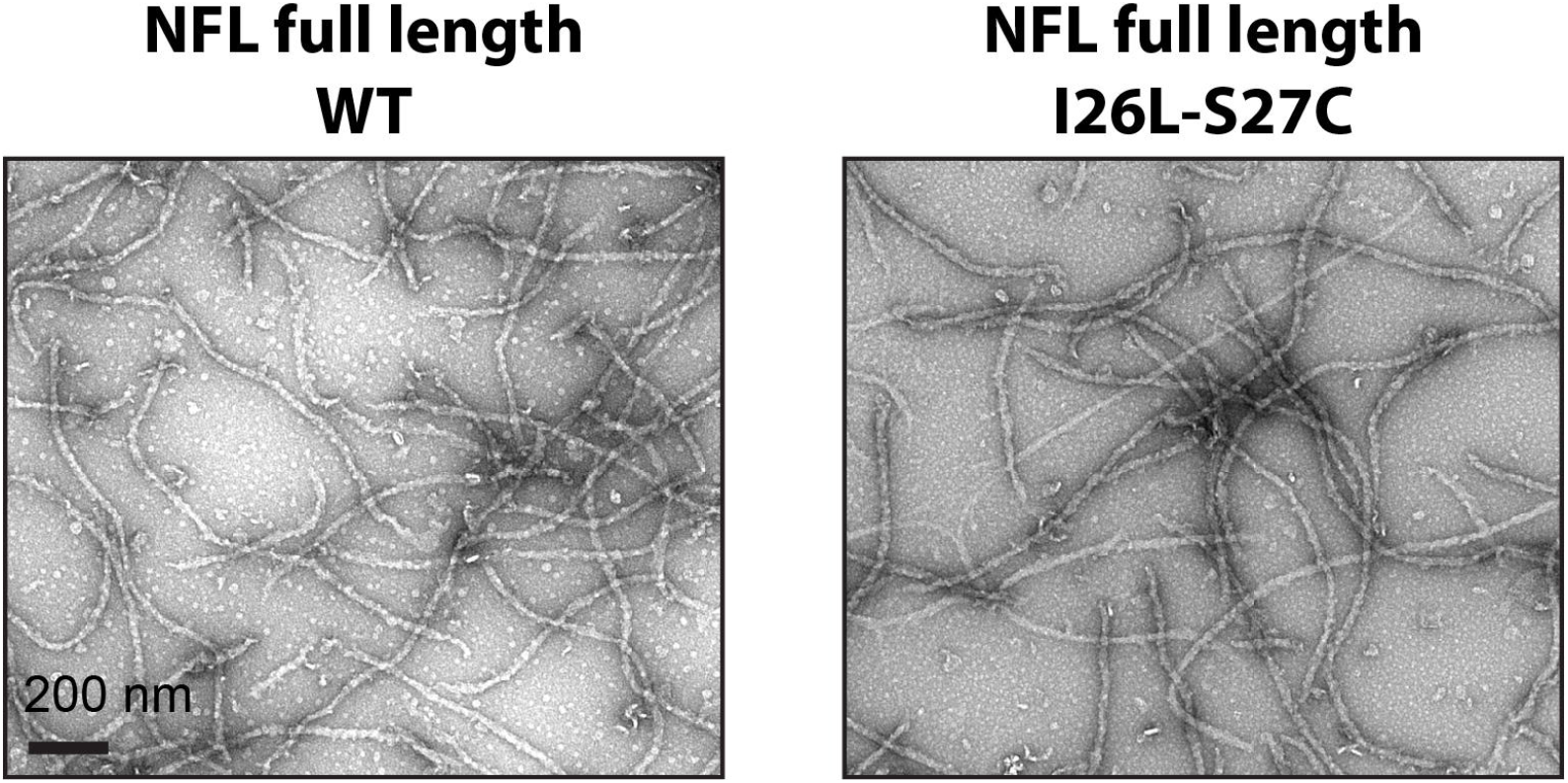

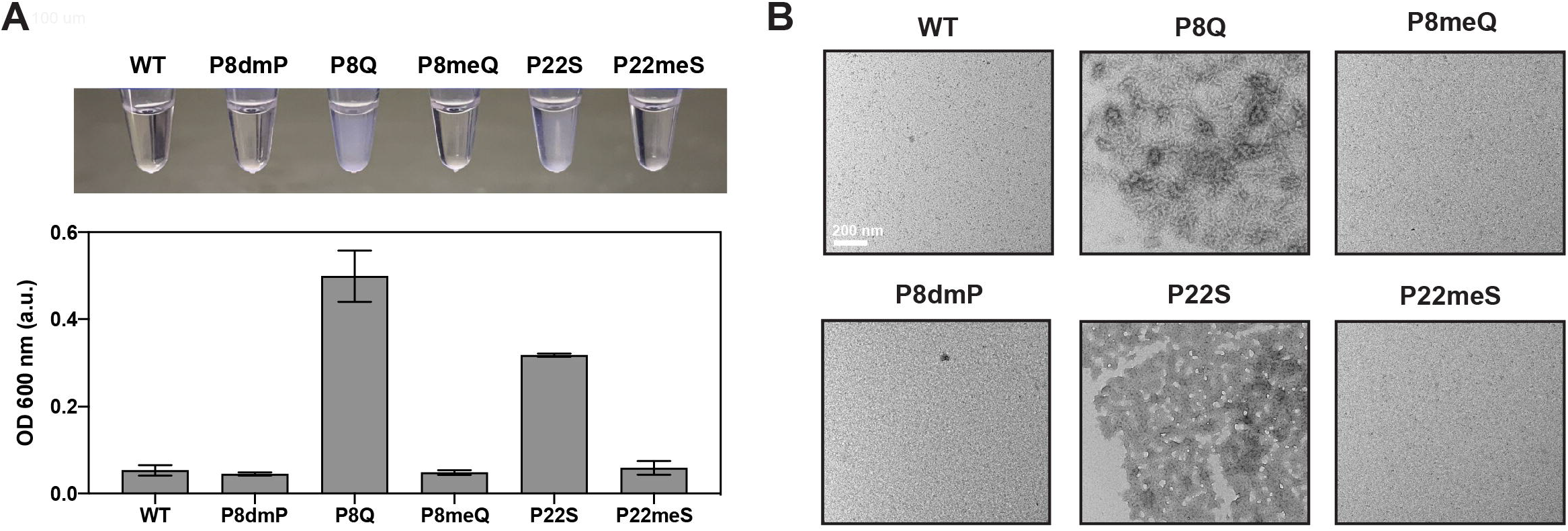

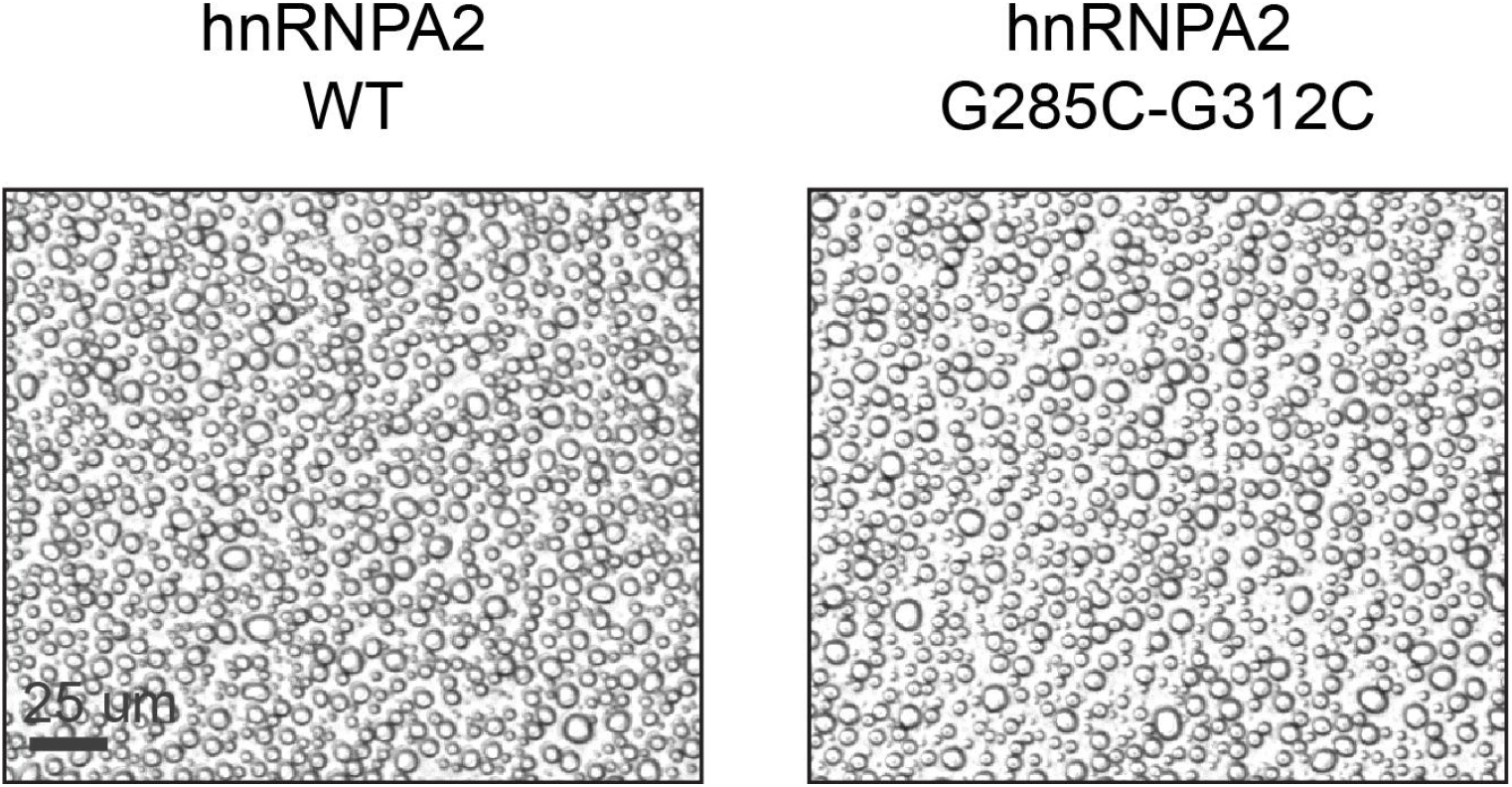

## Notes

### Competing Interest Statement

The authors have declared no competing interest.

