## Supplementary material for "Tracing the genetics of neurological disease to the mutation-directed addition of single hydrogen bonds": materials and methods, and supplementary figures

#### Molecular cloning

Standard gene cloning and mutagenesis strategies were used to generate expression constructs for all recombinant proteins. For native chemical ligation-compatible NFL fragments, an N-terminal 6×His-GFP tag was fused to constructs that would ultimately yield the head domain (residues 27-87) and full length (residues 27-543) constructs. A caspase 3 cleavage site (GDEVDC) was inserted between GFP and the NFL fragments such that digestion would yield NFL fragments bearing an N-terminal cysteine at amino acid position 27. These constructs were built into the pHis-parallel vector. Plasmid constructs for wild type full length NFL and its variants were prepared as previously described *(16)*.

The hnRNPA2-intein fusion fragment (hnRNPA2 residues 181-284 with a C-terminal GyrA intein fusion) was inserted in the pHis-parallel vector. A 6xHis tag was appended to C-terminus, and N-terminal 6×His tag was omitted during cloning. The N-terminal 6×His-tagged wild type hnRNPA2 LCD (residues 181-341) and its variants were made by inserting the LCD fragment into the pHis-parallel vector.

#### Recombinant NFL and hnRNPA2 fragment sequences

<u>hnRNPA2 (residues: 181-284)</u>
MQEVQSSRSGRGGNFGFGDSRGGGGNFGPGPGSNFRGGSDGYGSGRGFGDGYNGYGG
GPGGGNFGGSPGYGGGRGGYGGGGPGYGNQGGGYGGGYDNYGGGNYG-thioester

<u>NFL (S27C, residues 27-86, for head domain assembly)</u>
CSVRSGYSTARSAYSSYSAPVSSSLSVRRSYSSSSGSLMPSLENLDLSQVAAISNDLKSI

<u>NFL (S27C, residues 27-543, for full length assembly)</u>
CSVRSGYSTARSAYSSYSAPVSSSLSVRRSYSSSSGSLMPSLENLDLSQVAAISNDLKSIR
TQEKAQLQDLNDRFASFIERVHELEQQNKVLEAELLVLRQKHSEPSRFRALYEQEIRDLR
LAAEDATNEKQALQGEREGLEETLRNLQARYEEEVLSREDAEGRLMEARKGADEAALA
RAELEKRIDSLMDEISFLKKVHEEEIAELQAQIQYAQISVEMDVTKPDLSAALKDIRAQY
EKLAAKNMQNAEEWFKSRFTVLTESAAKNTDAVRAAKDEVSESRRLLKAKTLEIEACR
GMNEALEKQLQELEDKQNADISAMQDTINKLENELRTTKSEMARYLKEYQDLLNVKM
ALDIEIAAYRKLLEGEETRLSFTSVGSITSGYSQSSQVFGRSAYGGLQTSSYLMSTRSFPS
YYTSHVQEEQIEVEETIEAAKAEEAKDEPPSEGEAEEEEKDKEEAEEEEAAEEEEAAKEE
SEEAKEEEEGGEGEEGEETKEAEEEEKKVEGAGEEQAAKKKD

#### Protein expression and purification

All recombinant proteins used in this study were expressed in *E.coli* BL21 (DE3) cells. Cells were grown to an $OD_{600}$ of 0.6 in LB medium. Expression of NFL proteins was induced with 0.6 mM IPTG at 16° C for 16 h. Cells were harvested by centrifugation at 4,000 × g for 15 min. The hnRNPA2 LCD proteins were expressed via addition of 0.8 IPTG at 37° C and cells were harvested after 4 h.

For purification of NFL fragments, the cell pellets were resuspended in a lysis buffer containing 25 mM Tris-HCl (pH 7.5), 150 mM NaCl, 2 M Urea, 5 mM β-mercaptoethanol (β-ME), 20 mM imidazole, and disrupted by sonication. The cell lysates were clarified via centrifugation at 36,000 × g for 50 min and supernatants were applied to $Ni^{2+}$-NTA resin (Qiagen) and the column washed with the lysis buffer and eluted with lysis buffer supplemented with 300 mM imidazole. The eluted proteins were dialyzed against lysis buffer without imidazole for 3 h. The caspase-3 enzyme was then added to the protein solution for 2 h to cleave the GFP tag while generating the N-terminal cysteine. The protein solution was again passed over $Ni^{2+}$-NTA resin to capture the GFP fusion tag and any undigested protein. Flow through containing NFL fragments was collected, supplemented with 1% TFA, and purified via RP-HPLC (C4 preparative column). Purity was determined by analytical RP-HPLC and ESI-MS and pure fractions were pooled and lyophilized. Wild type full length NFL and its variants were purified as previously described *(16)*.

For purification of the hnRNPA2 LCD fragment, the cell pellet was resuspended in a lysis buffer containing 25 mM potassium phosphate (pH 7.2), 150 mM NaCl, 8 M Urea, 1 mM TCEP, 20 mM imidazole, and disrupted by sonication. The resulting cell lysates were centrifugated at 40,000 × g for 50 min. The supernatants were applied to $Ni^{2+}$-NTA resin and the column washed with lysis buffer and eluted with lysis buffer supplemented with 300 mM imidazole. The purified protein was refolded via overnight dialysis against a buffer containing 25 mM potassium phosphate (pH 7.2), 150 mM NaCl, 1 M Urea, 1 mM TCEP. The refolded protein was then centrifugated at 4,000 × g for 30 min to remove any precipitates, and the cleared supernatant was supplemented with 300 mM MES-Na and incubated at room temperature for 16 h. Following the GryA thiolysis reaction, the protein solution was spun at 4,000 × g for 30 min to pellet any precipitate. The precipitated protein was resolubilized by a buffer containing 25 mM potassium phosphate (pH 7.2), 150 mM NaCl, 6 M guanidine-HCl, 1 mM TCEP, and combined with supernatant fraction and loaded on $Ni^{2+}$-NTA resin. The flow through containing free hnRNPA2 LCD fragment was collected, supplemented with 1% TFA, and purified by RP-HPLC. Purity was determined by analytical RP-HPLC and ESI-MS and pure fractions were pooled and lyophilized.

**Peptide synthesis**

All fluorenylmethyloxycarbonyl (Fmoc)-protected amino acids were purchased from Oakwood Chemical or Combi-Blocks. Peptide synthesis resins (Trityl-OH ChemMatrix and Rink Amide ChemMatrix) were purchased from Biotage. All analytical reversed-phase HPLC (RP-HPLC) was performed on an Agilent 1260 series instrument equipped with a quaternary pump and an XBridge Peptide C18 or C4 column (5 µm, 4 × 150 mm; Waters) at a flow rate of 1 mL/min. Similarly, semi-preparative scale purifications were performed employing a XBridge Peptide C18 or C4 semi-preparative column (5 µm, 10 mm × 250 mm, Waters) at a flow rate of 4 mL/min. Preparative RP-HPLC was performed on an Agilent 1260 series instrument equipped with a preparatory pump and a XBridge Peptide C18 or C4 preparatory column (10 µM; 19 × 250 mm, Waters) at a flow rate of 20 mL/min. All instruments were equipped with a variable wavelength UV-detector. All RP–HPLC steps were performed using 0.1% (trifluoroacetic acid, TFA, Oakwood Chemical) in $H_2O$ (Solvent A) and 90% acetonitrile (Sigma-Aldrich), 0.1% TFA in $H_2O$ (Solvent B) as mobile phases. For LC/MS analysis, 0.1% formic acid (Sigma-Aldrich) was substituted for TFA in mobile phases. Mass analysis was carried out for each product on an LC/MSD (Agilent Technologies) equipped with a 300SB-C18 column (3.5 µM; 4.6 × 100 mm, Agilent Technologies) or a X500B QTOF (Sciex).

*Amidated peptides described in this study*
hnRNPA2 (306-341, S306C):
CGNFGGSRNMGGPYGGGNYGPGGSGGSGGYGGRSRY-$CONH_2$
tau-RD (295-311): DNIKHVPGGGSVQIVYK-$CONH_2$
tau-RD P301S (295-311, P301S): DNIKHVSGGGSVQIVYK-$CONH_2$
tau-RD P301meS (295-311, P301meS): DNIKHVmeSGGGSVQIVYK-$CONH_2$
tau-RD P301T (295-311, P301T): DNIKHVTGGGSVQIVYK-$CONH_2$
tau-RD P301meT (295-311, P301meT): DNIKHVmeTGGGSVQIVYK-$CONH_2$
tau-RD P301L (295-311, P301L): DNIKHVLGGGSVQIVYK-$CONH_2$
tau-RD P301meL (295-311, P301meL): DNIKHVmeLGGGSVQIVYK-$CONH_2$
tau-RD P301P-*cis* (295-311, P301P-*cis*): DNIKHVP*cis*GGGSVQIVYK-$CONH_2$
tau-RD P301P-*trans* (295-311, P301P-*trans*) DNIKHVP*trans*GGGSVQIVYK-$CONH_2$

where, me = $N^{\alpha}$-methyl

The above amidated peptides were synthesized via solid-phase peptide synthesis on a CEM Discover Microwave Peptide Synthesizer (Matthews, NC) using the Fmoc-protection strategy on Rink Amide-ChemMatrix resin (0.5 mmol/g). For coupling reactions, amino acids (5 eq) were activated with N,N-diisopropylcarbodiimide (DIC, 5 eq, Oakwood Chemical)/Oxyma (5 eq, Oakwood Chemical) and heated to 90 °C for 2 min while bubbling with $N_2$ in N,N dimethylformamide (DMF, Oakwood Chemical). Fmoc deprotection was carried out with 20% piperidine (Sigma-Aldrich) in DMF supplemented with 0.1 M 1-hydroxybenzotriazole hydrate (HOBt, Oakwood Chemical) at 90 °C for 1 min while bubbling with $N_2$. Cleavage from resin was performed with 92.5% TFA, 2.5% triisopropylsilane (TIS, SigmaAldrich), 2.5% 1,2-ethanedithiol (EDT, Sigma-Aldrich), and 2.5% $H_2O$ for 2 h at 25 °C on an end-over-end rotisserie. The crude peptide was then precipitated by the addition of a 10-fold volume of ice-cold ether and centrifuged at 4,000 RCF for 10 min at 4 °C. The pellet was resuspended in solvent A and purified via preparative C18 RP-HPLC. Fractions were analyzed on analytical C18 RP-HPLC and ESI-MS and those containing pure product (>95%) were pooled, lyophilized and stored at -80 °C.

*C-terminal SEA peptides described in this study*
NFL wild type (2-26, I26L): SSFSYEPYYSTSYKRRYVETPRVHL-SEA
NFL P8L (2-26, P8L, I26L): SSFSYELYYSTSYKRRYVETPRVHL-SEA
NFL P8meL (2-26, P8meL, I26L): SSFSYEmeLYYSTSYKRRYVETPRVHL-SEA
NFL P8Q (2-26, P8Q, I26L): SSFSYEQYYSTSYKRRYVETPRVHL-SEA
NFL P8meQ (2-26, P8meQ, I26L): SSFSYEmeQYYSTSYKRRYVETPRVHL-SEA
NFL P8R (2-26, P8R, I26L): SSFSYERYYSTSYKRRYVETPRVHL-SEA
NFL P8meR (2-26, P8meR, I26L): SSFSYEmeRYYSTSYKRRYVETPRVHL-SEA
NFL P8dmP (2-26, P8dmP, I26L): SSFSYEdmPYYSTSYKRRYVETPRVHL-SEA
NFL P8esL (2-26, P8esL, I26L): SSFSYEesLYYSTSYKRRYVETPRVHL-SEA
NFL P22R (2-26, P22R, I26L): SSFSYEPYYSTSYKRRYVETRRVHL-SEA
NFL P22meR (2-26, P22meR, I26L): SSFSYEPYYSTSYKRRYVETmeRRVHL-SEA

NFL P22S (2-26, P22S, I26L): SSFSYEPYYSTSYKRRYVETSRVHL-SEA
NFL P22meS (2-26, P22meS, I26L): SSFSYEPYYSTSYKRRYVETmeSRVHL-SEA
NFL P22T (2-26, P22T, I26L): SSFSYEPYYSTSYKRRYVETTRVHL-SEA
NFL P22meT (2-26, P22meT, I26L): SSFSYEPYYSTSYKRRYVETmeTRVHL-SEA
hnRNP A2 wild type (285-305, A285Thz): Thz-GNYNDFGNYNQQPSNYGPMK-SEA
hnRNP A2 P298L (285-305, P298L, A285Thz): Thz-GNYNDFGNYNQQLSNYGPMK-SEA
hnRNP A2 P298meL (285-305, P298meP, A285Thz): Thz-GNYNDFGNYNQQmeLSNYGPMK-SEA

where, SEA = bis(2-sulfanylethyl)amido group, dmP = L-5,5-Dimethylproline, me = $N^{\alpha}$-methyl, es = ester, Thz = thiazolidine

All peptides containing a C-terminal SEA moiety were synthesized similarly to the amidated peptides described above with the following modifications. For manual loading of the first amino acid to SEA resin (0.16 mmol/g; Iris Biotech), Fmoc-lysine (10 eq) or Fmoc-leucine (10 eq), HATU (10 eq, Oakwood Chemical), and DIPEA (30 eq) were mixed in DMF and the resin was bubbled with $N_2$ for 1 h. This step was repeated with fresh reagents to ensure complete loading. The resin was then washed with DMF and bubbled in acetic anhydride:DIPEA (20 eq:40 eq) in DMF for 20 min to quench unreacted sites. Subsequent peptide synthesis reactions were performed on an automated, microwave synthesizer as described for amidated peptides. Resin cleavage was performed with 95% TFA, 2.5% TIS, and 2.5% $H_2O$ for 2 h at 25 °C on an end-over-end rotisserie. Purification of SEA peptides were performed as described for amidated peptides.

For peptide chain elongation from the dmP building block, resin was removed from the synthesizer and manual coupling was performed with Fmoc-5,5-dmP-OH (6 eq, Iris Biotech), PyAOP (6 eq, Oakwood Chemical) and DIPEA (12 eq, SigmaAldrich) for 1 hour at 50 °C. Coupling was repeated with fresh reagents to maximize yield. Resin was acetyl capped (acetic anhydride:DIPEA, 20 eq:40 eq in DMF) to quench unreacted sites and placed back on automated synthesizer to complete synthesis.

The L-leucic acid building block was manually coupled to resin in a reaction containing L-leucic acid (5 eq, Combi-Blocks), DIC (5 eq) and oxyma (5 eq) in DMF for 30 min at 20 °C. Coupling was repeated twice with fresh reagents to maximize yield. For peptide chain elongation from the L-leucic acid building block, a reaction containing amino acid (5 eq, Fmoc-Glu(OtBu)-OH), MSNT (5 eq, SigmaAldrich), and 1-methylimidazole (3.75 eq, SigmaAldrich) in DCM was purged with $N_2$ and incubated for 2 h on an end-over-end rotisserie. Coupling was repeated thrice with fresh reagents to maximize yield. Following final reaction, resin was placed back on automated synthesizer to complete synthesis.

### Assembly of semi-synthetic NFL constructs

To assemble NFL head domains (residues 2-86), native chemical ligation reactions were performed by combining 5 mM peptide bearing C-terminal SEA moieties (NFL residues 2-26; wt, P8Q, PmeQ, P8dmP, P22S or P2meS) with 1 mM of the NFL head domain fragment bearing an N-terminal cysteine (S27C, residues 27-86) in a degassed buffer of 6 M guanidine-HCl, 0.1 M sodium phosphate, 20 mM TCEP, 200 mM MES-Na (Sigma-Aldrich), and 150 mM 2,2,2-

trifluoroethanethiol (TFET, Sigma-Aldrich). Reaction pH was adjusted to 7.0 and incubated at 37 °C for 16 h. Reaction progress was monitored via RP-HPLC and ESI-MS analysis. Ligation products were purified on a preparative C18 RP-HPLC column and fractions were analyzed via analytical RP-HPLC and ESI-MS. Fractions containing target mass were pooled, lyophilized, and stored at -80 °C.

To assemble full length NFL (2-543), native chemical ligation reactions were performed by combining peptides bearing C-terminal SEA moieties (NFL 2-26, wt, P8L, P8esL, P8meL, P8Q, P8meQ, P8R, P8meR, P8dmP, P22S, P22meS, P22R, P22meR, P22T, P22meT) with the NFL truncated fragment bearing N-terminal cysteine (S27C, residues 27-543). Ligation reactions were performed as described for isolated NFL head domain assembly.

**Assembly of semi-synthetic hnRNPA2 LCD constructs**

To assemble the hnRNPA2 LCD (residues 181-341) a three-piece native chemical ligation strategy was devised. In Ligation 1, peptides bearing an N-terminal Thz and C-terminal SEA moiety (hnRNPA2 residues 285-305, hnRNPA2 residues 285-305 with P298L mutation, or hnRNPA2 residues 285-305 with P298meL mutation) were combined with the hnRNPA2 C-terminal peptide (residues 306-341, S306C). Reactions included 1 mM SEA peptide, 0.5 mM C-amidated peptide, 200 mM MES-Na, 20 mM TCEP, and 150 mM TFET in a degassed buffer of 6 M guanidine-HCl and 0.1 M sodium phosphate at pH 7.0. Reactions were incubated at 37 °C for 16 h and progress was monitored via RP-HPLC and ESI-MS analysis. Upon completion of the reaction, Thz deprotection was initiated by addition of 200 mM O-methylhydroxylamine-HCl and 20 mM TCEP in a degassed buffer of 6 M guanidine-HCl and 0.1 M sodium phosphate. Reactions were adjusted to pH 4.0 and incubated at 37 °C for 4 h. Products were purified on a preparative C18 RP-HPLC column and analyzed via analytical RP-HPLC and ESI-MS. Fractions containing target mass were pooled, lyophilized and stored at -80 °C. These products (wild type, P298L and P298meL) are referred to as hnRNPA2 Piece 2+3.

For Ligation 2, the hnRNPA2 Piece 2+3 constructs, each bearing a free N-terminal cysteine, were combined with the recombinant hnRNPA2 thioester construct (residues 181-284). These reactions included 1 mM Piece 2+3, 0.5 mM recombinant hnRNPA2 thioester fragment, 20 mM TCEP, and 150 mM TFET in a degassed buffer of 6 M guanidine-HCl and 0.1 M sodium phosphate at pH 7.0. Reactions were incubated at 37 °C for 16 h and progress was monitored via RP-HPLC and ESI-MS analysis. Upon completion of the reactions, the ligation products were purified on a semi-preparative C18 RP-HPLC column and fractions were characterized via RP-HPLC and ESI-MS. Pure fractions were pooled, lyophilized and stored at -80 °C.

**Phenotypic analysis of NFL filament assembly, NFL N-terminal peptide solubility and head domain polymer formation**

NFL filament assembly was performed as previously described.

To test NFL peptide solubility, the lyophilized peptides were first completely dissolved with neat TFA, which was then evaporated under a gentle stream of $N_2$. Residual TFA was removed via lyophilization for 30 min. This step is important to ensure peptides are completely disaggregated prior to Thioflavin-T analysis. Lyophilized peptides were then dissolved in a buffer containing

25 mM Tris-HCl (pH 7.5), 150 mM NaCl, 5 mM β-ME, 6 M guanidine-HCl and diluted into PBS supplemented with 20 μM Thioflavin-T (final peptide concentration = 200 μM). Solutions were next loaded into a 96-well clear bottom plate and incubated at 4 °C for 5 min. Precipitation was also imaged on a ZOE Fluorescent Cell Imager (Bio-Rad).

To test NFL head domain polymer formation, each variant was dissolved in a buffer containing 25 mM Tris-HCl (pH 7.5), 150 mM NaCl, 5 mM β-ME, and 8 M urea, and dialyzed against an identical buffer with a reduced urea concentration (3 M urea). Sample turbidity was measured by absorbance at 600 nm and head domain polymers were confirmed by negative-staining EM.

**Thioflavin-T assay for tau peptides**

Lyophilized tau peptides were first completely dissolved with neat TFA, which was then evaporated under a gentle stream of $N_2$. Residual TFA was removed via lyophilization for 30 min *(30)*. Next, peptides were dissolved in a buffer containing 25 mM Tris-HCl (pH 7.5), 150 mM NaCl, 6 M guanidine-HCl and diluted into PBS supplemented with 20 μM Thioflavin-T (final peptide concentration = 100 μM). Solutions were loaded into a 96-well fluorescence assay plate, sealed (Constar 6570 sealing tape) and Thioflavin-T fluorescence was monitored for 99 h (Cytation 5 plate reader, BioTek). Five repeats were performed for all samples.

**Phenotypic analysis of tau peptides in tau biosensor cell line**

The tau biosensor cell line was purchased from ATCC and cultured in a Dulbecco's modified Eagle's medium (DMEM) supplemented with 10% fetal bovine serum. Tau biosensor cells were seeded in a 96-well tissue culture plate a day prior to tau peptide transfection.
Tau peptides were disaggregated, diluted in PBS to 300 μM, and sonicated in a water bath sonicator (BRANSON 2800) for 2 min. Tau peptide solution was mixed with same volume of a lipofectamine 3000-OptiMEM solution incubated for 30 min at room temperature. Ten μl of tau peptide transfection mixture was added to 90 μl of tau biosensor cells to reach a final peptide concentration of 15 μM in medium. The cells were imaged by EVOS FL Fluorescence Microscope 48-72 h post transfection.

**Phenotypic analysis of hnRNPA2 LCD phase-separated droplets**

hnRNPA2 LCD proteins were dissolved in a buffer containing 25 mM Tris-HCl (pH 7.5), 150 mM NaCl, 6 M guanidine-HCl. hnRNPA2 LCD phase-separated droplet formation was induced by diluting protein in either a buffer containing 25 mM Tris (pH 7.5), 150 mM NaCl, 5 mM β-ME, or a buffer containing 50 mM 2-(N-Morpholino)ethanesulfonic acid (pH 5.5), 100 mM NaCl, 5 mM β-ME to a final concentration of 15 μM. Identical results were observed for the wild type hnRNPA2 LCD and its variants in these two buffers. However, the average size of hnRNPA2 droplets increased in the pH 5.5 buffer and these conditions were used for image analysis via the ZOE Fluorescent Cell Imager.

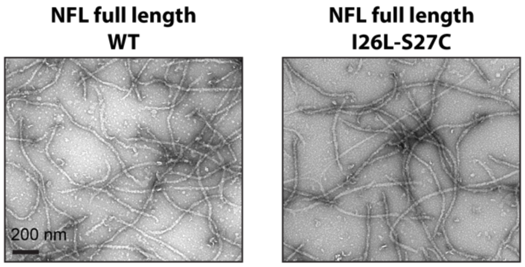


**Fig. S1. The I26L/S27C variant of NFL forms intermediate filaments indistinguishable from the native protein.**

The native chemical ligation approach used to prepare semi-synthetic derivatives of NFL led to the deposition of two missense variations at the junction site wherein synthetic peptides were fused to the remainder of the native NFL protein. Isoleucine residue 26 of the native protein was replaced by leucine, and serine residue 27 was replaced by cysteine. Recombinant NFL proteins bearing either the native NFL sequence or the I26L/S27C double variant were expressed in bacteria, purified and incubated under conditions receptive to the assembly of intermediate filaments (Materials and Methods). Both proteins assembled into indistinguishable, homogeneous filaments as revealed by transmission electron microscopy. Scale bar = 200 nm.

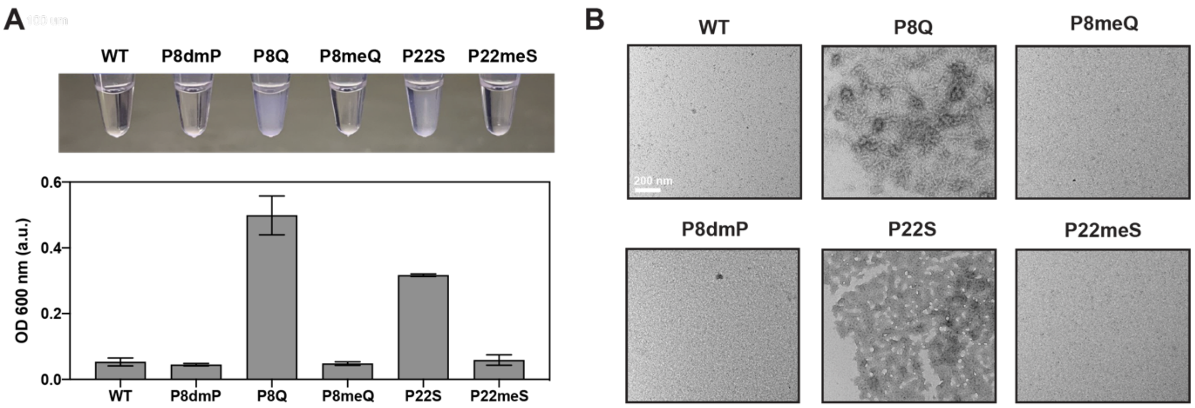


**Fig. S2. Restorative effects of eliminating Pauling hydrogen bonding by methyl-capping the peptide backbone nitrogen atom of the P8Q and P22S mutational variants within the NFL head domain.**

Peptides were synthesized corresponding to: (i) the native sequence of the first twenty five residues of NFL; (ii) a variant replacing proline residue 8 with dmP; (iii) variants replacing proline 8 with glutamine, or proline 22 with serine; or (iv) variants in which the peptide backbone nitrogens associated with glutamine 8, or serine 22 were methyl-capped. Native chemical ligation was used to conjugate each peptide to the remainder of the NFL head domain (Materials and Methods). Semi-synthetic head domains were purified and incubated under conditions known to differentiate the solubility of the native head domain relative to variants carrying individual CMT mutations. The native protein (WT) and the variant carrying dmP in place of proline residue 8 (P8dmP) were soluble under such conditions. The P8Q and P22S variants were insoluble, and methyl-capping of the peptide backbone nitrogen atoms associated with glutamine 8 and serine 22 restored solubility **(Panel A)**. Each sample was evaluated by transmission electron microscopy, revealing precipitates in the head domain samples containing the CMT variants **(Panel B)**. Scale bar = 200 nm.

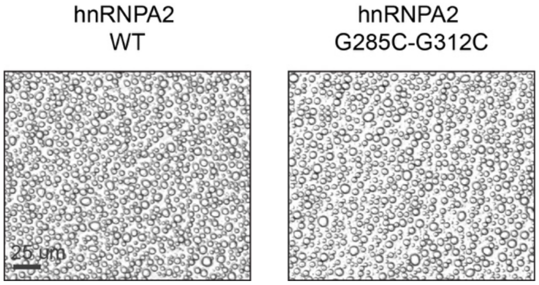


**Fig. S3. The G285C/G312C variant of the hnRNPA2 low complexity domain forms liquid-like droplets indistinguishable from droplets formed by native protein.**

The three-piece native chemical ligation approach used to prepare semi-synthetic derivatives of the hnRNPA2 LC domain led to the deposition of a missense mutation at both junction sites wherein the synthetic peptide was fused with to the remaining N- and C-terminal regions of the protein (Materials and Methods). Glycine residue 285 was changed to cysteine, and glycine residue 312 was also changed to cysteine. Proteins corresponding to the native hnRNPA2 LC domain and the G286C/G312C variant were expressed in bacteria, purified and subjected to assays of phase separation. The two proteins formed indistinguishable, spherical liquid-like droplets. Scale bar = 25 µm.

**Data S1. (Separate file)**

Mass spectrometry characterization of semi-synthetic native neurofilament light (NFL), semi-synthetic native hnRNPA2 low complexity domain, native tau peptide, and variants of all the three above proteins.
